# Lung microenvironments harbor *Mycobacterium tuberculosis* phenotypes with distinct treatment responses

**DOI:** 10.1101/2023.02.03.527062

**Authors:** Nicholas D. Walter, Jackie P. Ernest, Christian Dide-Agossou, Allison A. Bauman, Michelle E. Ramey, Karen Rossmassler, Lisa M. Massoudi, Samantha Pauly, Reem Al Mubarak, Martin I. Voskuil, Firat Kaya, Jansy P. Sarathy, Matthew D. Zimmerman, Véronique Dartois, Brendan K. Podell, Radojka M. Savic, Gregory T. Robertson

**Affiliations:** Rocky Mountain Regional VA Medical Center, Aurora, CO, USA; Division of Pulmonary Sciences and Critical Care Medicine, University of Colorado Anschutz Medical Campus, Aurora, CO, USA; Consortium for Applied Microbial Metrics, Aurora, CO, USA; Department of Bioengineering and Therapeutic Sciences, University of California San Francisco, San Francisco, CA, USA; Mycobacteria Research Laboratories, Department of Microbiology, Immunology and Pathology, Colorado State University, Fort Collins, Colorado, USA; Department of Immunology and Microbiology, University of Colorado Anschutz Medical Campus, Aurora, CO, USA; Center for Discovery and Innovation, Hackensack Meridian Health, Nutley, New Jersey, USA

## Abstract

Tuberculosis lung lesions are complex and harbor heterogeneous microenvironments that influence antibiotic effectiveness. Major strides have been made recently in understanding drug pharmacokinetics in pulmonary lesions, but the bacterial phenotypes that arise under these conditions and their contribution to drug tolerance is poorly understood. A pharmacodynamic marker called the RS ratio quantifies ongoing rRNA synthesis based on the abundance of newly-synthesized precursor rRNA relative to mature structural rRNA. Application of the RS ratio in the C3HeB/FeJ mouse model demonstrated that *Mycobacterium tuberculosis* populations residing in different tissue microenvironments are phenotypically distinct and respond differently to drug treatment with rifampin, isoniazid or bedaquiline. This work provides a foundational basis required to address how anatomic and pathologic microenvironmental niches may contribute to the long treatment duration and drug tolerance during treatment of human tuberculosis.

## INTRODUCTION

To address the ongoing global tuberculosis (TB) epidemic, there is a critical need for new shorter, more effective combination antibiotic regimens. Current standard treatments require six months for treatment of drug-susceptible TB and may take years for certain drug-resistant forms of TB. Development of novel regimens that cure TB more rapidly will require better understanding of how drugs and regimens affect treatment-refractory bacterial populations.

One reason that TB demands prolonged therapy is the complexity and heterogeneity of TB lung lesions.^1–3^ Lesional microenvironments influence antibiotic effectiveness in two distinct ways. First, the variable architecture and composition of lesions, including differences in vascularization, fibrosis, inflammatory and immune cell infiltration and activation, and caseum formation, affect drug penetration and retention, resulting in drug- and lesion-specific pharmacokinetic (PK) profiles.^2, 4–6^ Second, variable physiochemical properties, including levels of O_2_, CO_2_, H^+^, carbon and nitrogen sources, micronutrients, and host immune milieu elicit differing metabolic and stress responses in *Mycobacterium tuberculosis (Mtb*), resulting in phenotypically distinct bacterial populations in lesions that tolerate drug exposure differently. For example, *Mtb* in rabbit caseum are largely non-replicating and exhibit extreme drug tolerance to many drugs.^3^ A similar dependence of antibiotic effectiveness on lesional phenotypes has been observed in non-human primates and marmosets (reviewed in^2^). Antibiotic effectiveness within a lesion depends on both PK (*i.e*., drugs must reach the pathogen at therapeutically relevant concentrations) and on the *Mtb* phenotype present (*i.e*., drugs must have activity agains the lesionspecific phenotype).^7^ While important strides have recently been made in understanding PK distribution of antituberculosis agents into pulmonary lesions in diverse animal models of TB,^6,8,9^ there remains limited knowledge of lesion *Mtb* phenotypes and how these distinct phenotypes may respond differently to drug exposure.^10–14^

Drug effects have traditionally been assessed based on colony forming units (CFU) which enumerate the burden of bacteria capable of growth on agar. We recently described the RS ratio,^15^ an alternative pharmacodynamic (PD) marker that is conceptually distinct from measures of bacterial burden. The RS ratio quantifies a key bacterial cellular process: ongoing rRNA synthesis. rRNA synthesis is fundamentally linked with bacterial replication.^16,17^ In the absence of drug treatment, the RS ratio is a proxy for *Mtb* replication.^15^ We have previously shown that the potency of a regimen in suppressing the RS ratio is associated with shortening time to non-relapsing cure in mice.^15^

Here we used both CFU and the RS ratio to evaluate the effect of three clinically relevant antibiotics in tissue microenvironments of the C3HeB/FeJ (Kramnik) mouse. We demonstrate that, in the absence of treatment, the *Mtb* populations of caseum, airway, remaining lung and spleen exhibit distinct phenotypes that have a spectrum of ongoing rRNA synthesis. Use of the RS ratio reveals that drugs with diverse mechanisms of action differently affect *Mtb* microenvironment-specific phenotypes. Collectively, this work demonstrates that advancing beyond enumeration of bacterial burden to consider cellular activity of unique *Mtb* phenotypes provides insight into drug effect in lesions and how microenvironmental niches contribute to the long treatment durations required to cure TB.

## RESULTS

### Pathology of C3HeB/FeJ mouse microenvironments

After 12-weeks of infection, C3HeB/FeJ mice had diverse and complex pathology that enabled evaluation of four microenvironments analgous to lesions observed in human TB.^2,18,19^ Discrete well-circumscribed granulomas classified as Type I lesions^2,18,19^ in which the central caseum is bounded by a peripheral rim of enlarged, vacuolated macrophages and an outer ring of fibrosis and compressed alveolar lung tissue were surgically incised to extract caseum (**Fig. 1a-b**). The *Mtb* population in caseum is predominantly extracelluar.^18,19^

**Fig 1.**
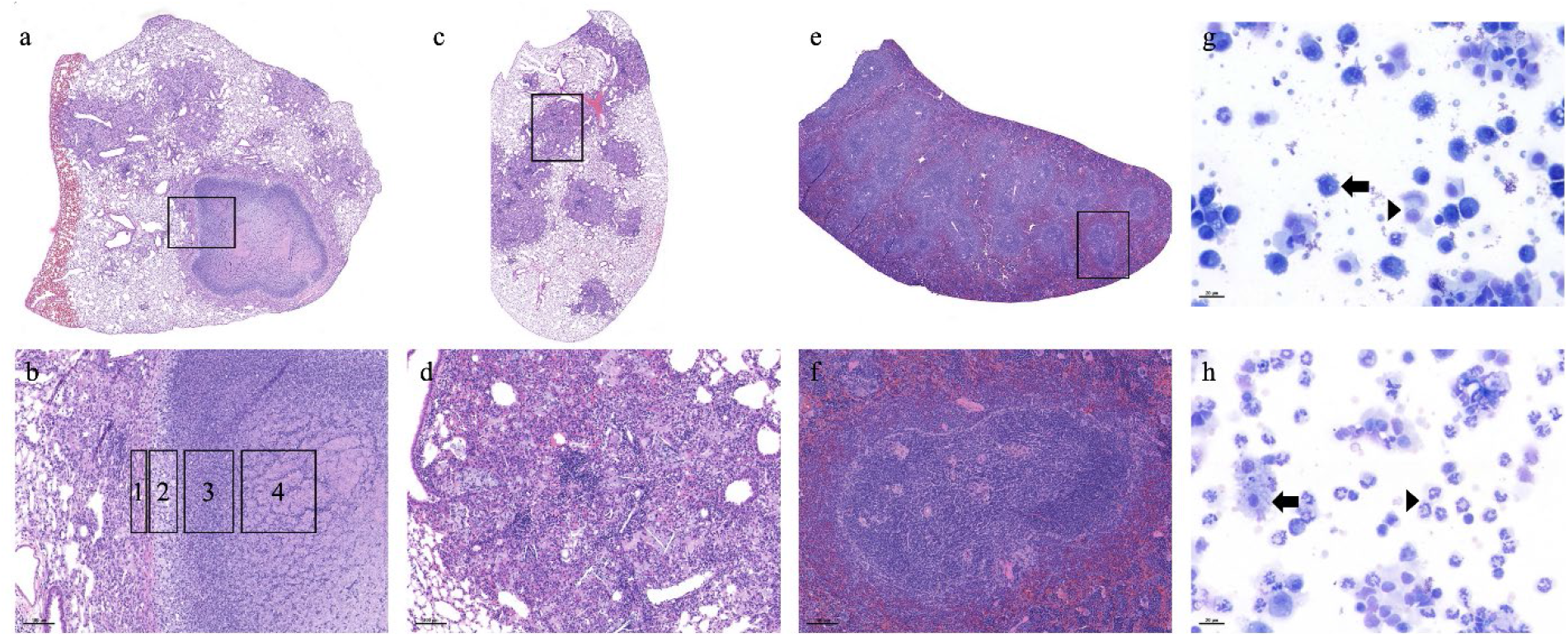
Pathology of four key microenvironments of the C3HeB/FeJ mouse evaluated in this manuscript. The caseum is the central component of the highly organized, caseous necrotic granuloma previously described as a Type I lesion (A and inset shown in B),^2,18,19^ Type I lesions are bordered by a rim of fibrosis and compressed lung tissue with interstitial inflammation (B, zone 1). Immediately central to the rim is a thin layer of vacuolated macrophages with high intracellular *Mtb* burden (B, zone 2). The caseum sampled in this study consists of an outer zone of neutrophil-dominated inflammation and inner zone of cellular debris (B, zone 3 and 4). The caseum is hypoxic, exhibits a near neutral pH, and contains a high burden of extracellular *Mtb*. Remaining lung is the lung parenchyma remaining after excision of visible Type 1 lesions. Remaining lung consists of type III lesions (C and inset shown in D), which are cellular non-necrotizing lesions with a relatively low burden of *Mtb* that is intracellular. Spleen (E and inset shown in F) has minimal infection-associated pathology with small aggregates of macrophages contained within the white pulp regions (arrow). Airway sampled via bronchoalveolar lavage consists of alveolar macrophages (arrow) and shed airway epithelial cells (arrowhead) in the uninfected animal (G), which shifts to neutrophil dominated inflammation (arrowhead) among fewer macrophages (arrow) in the infected lung prior to initiation of antimicrobial treatment.

The second microenvironment was remaining lung after removal of Type I lesions. Remaining lung was dominated by non-necrotic cellular Type III lesions consisting of an admixture of macrophages and lymphocytes with few neutrophils (Fig. 1c-d). The *Mtb* population of remaining lung is predominantly intracellular within macrophages.^18,19^

The third microenvironment was the airway which was sampled via bronchoalveolar lavage. In contrast to uninfected mice in which airway lavage is dominated by alveolar macrophages and epithelial cells (**Fig 1g**), lavage from untreated *Mtb-*infected mice revealed higher proportions of neutrophils and fewer macrophages (**Fig 1h**).

The fourth microenvironment was the spleen. Although spleen contains culturable *Mtb*, the pathological response is limited, consisting of microscopic aggregates of macrophages localized within lymphoid white pulp compartments (**Fig. 1e-f**).

### CFU burden and rRNA synthesis prior to drug treatment in key microenvironmental niches

Prior to drug treatment, CFU per gram was highest in caseum with levels at least 3 log10 CFU higher than any other microenvironment (minimum *P-*value <0.00001) (**Fig 2a**, Supplemental Table 1). CFU in remaining lung was lower than caseum but 0.78 log10 CFU higher than spleen (P=0.02). The RS ratio identified a range of *Mtb* rRNA synthesis rates from lowest in caseum, intermediate in remaining lung and spleen and highest in airway, indicating distinct microenvironment-specific phenotypes (**Fig 2b**). Differences between the RS ratio in each microenvironment were statistically significant (Supplemental Table 1).

**Fig 2.**
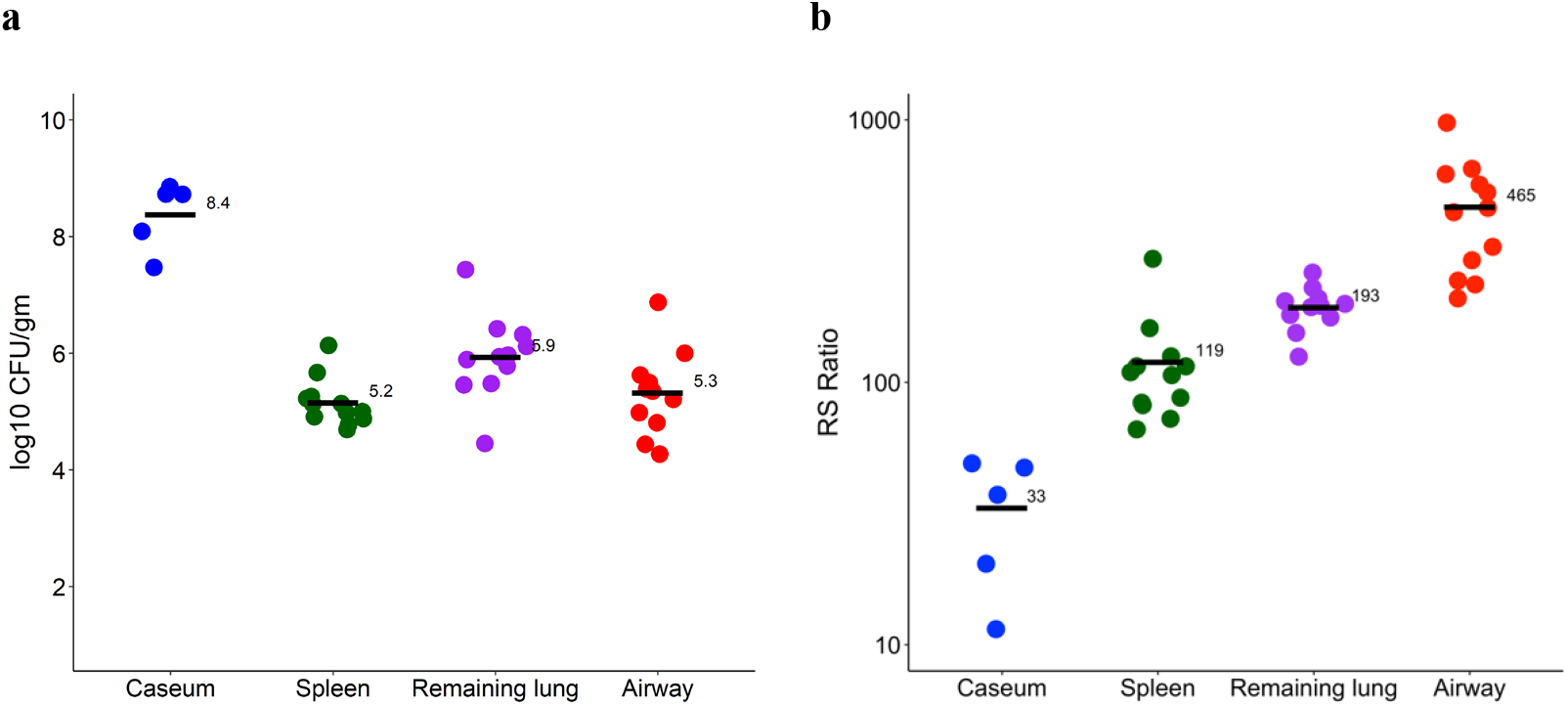
CFU (**a**) and RS ratio (**b**) in key microenvironments of the C3HeB/FeJ mouse prior to drug treatment. Circles represent values from individual tissue samples and horizontal bars indicate group means. Caseum, spleen, remaining lung and airway are represented in blue, green, purple and red, respectively.

Higher CFU was not associated with higher RS ratios. For example, the microenvironment with the highest *Mtb* burden (caseum) had the lowest RS ratio, consistent with a large *Mtb* population and low ongoing rRNA synthesis, suggestive of a slowed bacterial replication rate in caseum. The less concentrated *Mtb* population in remaining lung had an RS ratio 6-fold higher than caseum, CFU was similar in spleen and airway but airway had a significantly higher RS ratio (*P-*value <0.00001), suggesting an increased rate of bacterial replication in the oxygen-rich airway. As an additional point of reference, we assayed the RS ratio in the artificial *ex vivo* caseum surrogate model used to assess nonspecific caseum drug binding^20,21^ or drug activity against *Mtb* phenotypes that arise after incubation in lipid-rich caseum surrogate.^22^ The median RS ratio values were 45 in *ex vivo* caseum surrogate and 33 in caseum from C3HeB/FeJ mice, suggesting the *Mtb* phenotypes are in similar states of rRNA synthesis and replication.

### Effect of different drugs in specific microenvironments

Following 2.5 weeks of daily treatment, we observed that isoniazid (INH), rifampin (RIF10 or RIF30) and bedaquiline (BDQ) had differing effects within each individual microenvironment (**Fig 3**, Supplemental Table 2).

**Fig 3.**
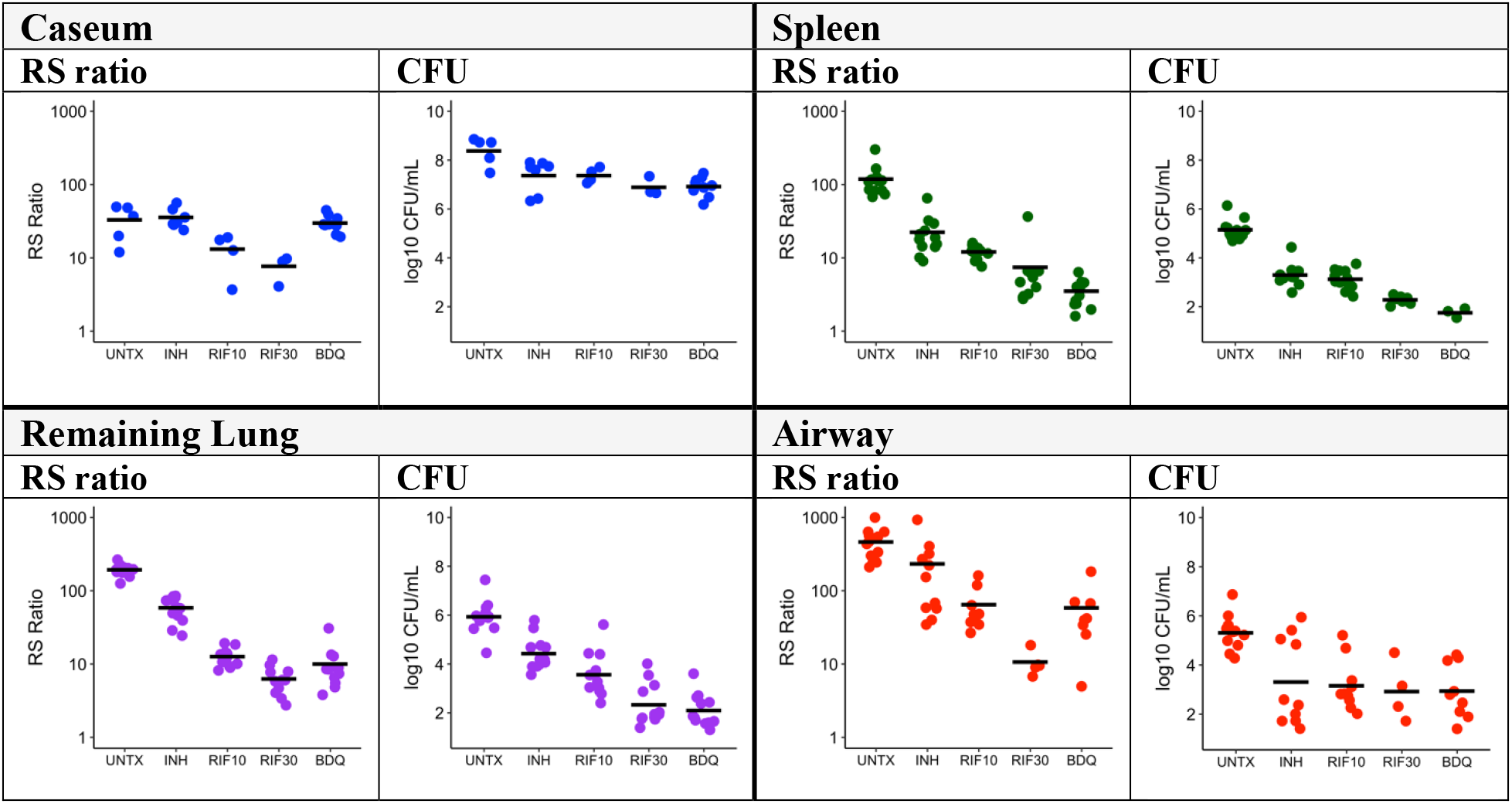
Effect of different drugs and doses on CFU and the RS ratio within specific microenvironments of the C3HeB/FeJ mouse. The RS ratio and CFU are shown for caseum (**a-b**), spleen (**c-d**), remaining lung (**e-f**) and airway (**g-h**). Dots indicate individual mice. Horizontal lines indicate group means. Caseum, spleen, remaining lung and airway are represented in blue, green, purple and red, respectively.

### Drug effect in caseum

In caseum, INH and BDQ had no effect on the RS ratio (*P*=0.97 and 0.99 relative to control, respectively). By contrast, RIF10 and RIF30 reduced the RS ratio 2.6-fold and 4.1-fold (*P*=0.04 and 0.003 relative to control, respectively). RIF30 reduced the RS ratio significantly more than INH or BDQ (*P*=0.0004 and 0.001, respectively). In caseum, CFU did not identify significant differences between drugs. All drugs and doses reduced CFU in caseum by approximately 1 log_10_ relative to the untreated controls.

### Drug effect in spleen

In the spleen, INH and RIF10 reduced the RS ratio 5.6-fold and 9.2-fold relative to control (minimum *P*<0.00001). RIF30 and BDQ both had a significantly greater effect, reducing the RS ratio at least 20-fold relative to control (minimum *P*<0.00001) and significantly more than INH (minimum *P*<0.00001). In contrast to caseum where BDQ had no effect on the RS ratio, BDQ was as effective as RIF30 in the spleen (*P*=0.1). CFU did identify significant differences between drugs in the spleen. Specifically, RIF30 or BDQ reduced CFU burdens in spleen more than INH (minimum *P*<0.00001) or RIF10 (minimum *P*=0.00005).

### Drug effect in remaining lung

In remaining lung, INH reduced the RS ratio 3.5-fold relative to control (*P*<0.00001). RIF10, RIF30 and BDQ had significantly greater effect, reducing the RS ratio at least 15-fold relative to control (minimum *P*<0.00001) and significantly more than INH (minimum *P*=<0.00001). Similar to results in the spleen, BDQ was as effective as RIF30 based on RS ratio in remaining lung (*P*=0.12). CFU revealed significant differences between drugs in remaining lung. Specifically, RIF30 or BDQ reduced CFU burdens more than INH (minimum *P*<0.00001) or RIF10 (minimum *P*=0.002).

### Drug effect in airway

In the airway, INH reduced the RS ratio 3-fold relative to control (P=0.01). RIF10, RIF30 and BDQ had significantly greater effect, reducing the RS ratio at least 8-fold relative to control (minimum *P*<0.00001). RIF30 had the greatest effect in airway, reducing the RS ratio significantly more than RIF10 or BDQ (minimum *P*=0.03). In airway, CFU did not identify significant differences between drugs. All drugs and doses reduced CFU by approximately 2 log10 relative to untreated controls in airway.

### Effects of individual drugs across different microenvironments

Using the RS ratio to evaluate individual drugs in different microenvironments revealed drug-specific changes (**Fig 4**, Supplemental Tables 3, 4). We evaluated both the change in RS ratio relative to control and the absolute RS ratio value following treatment.

**Fig 4.**
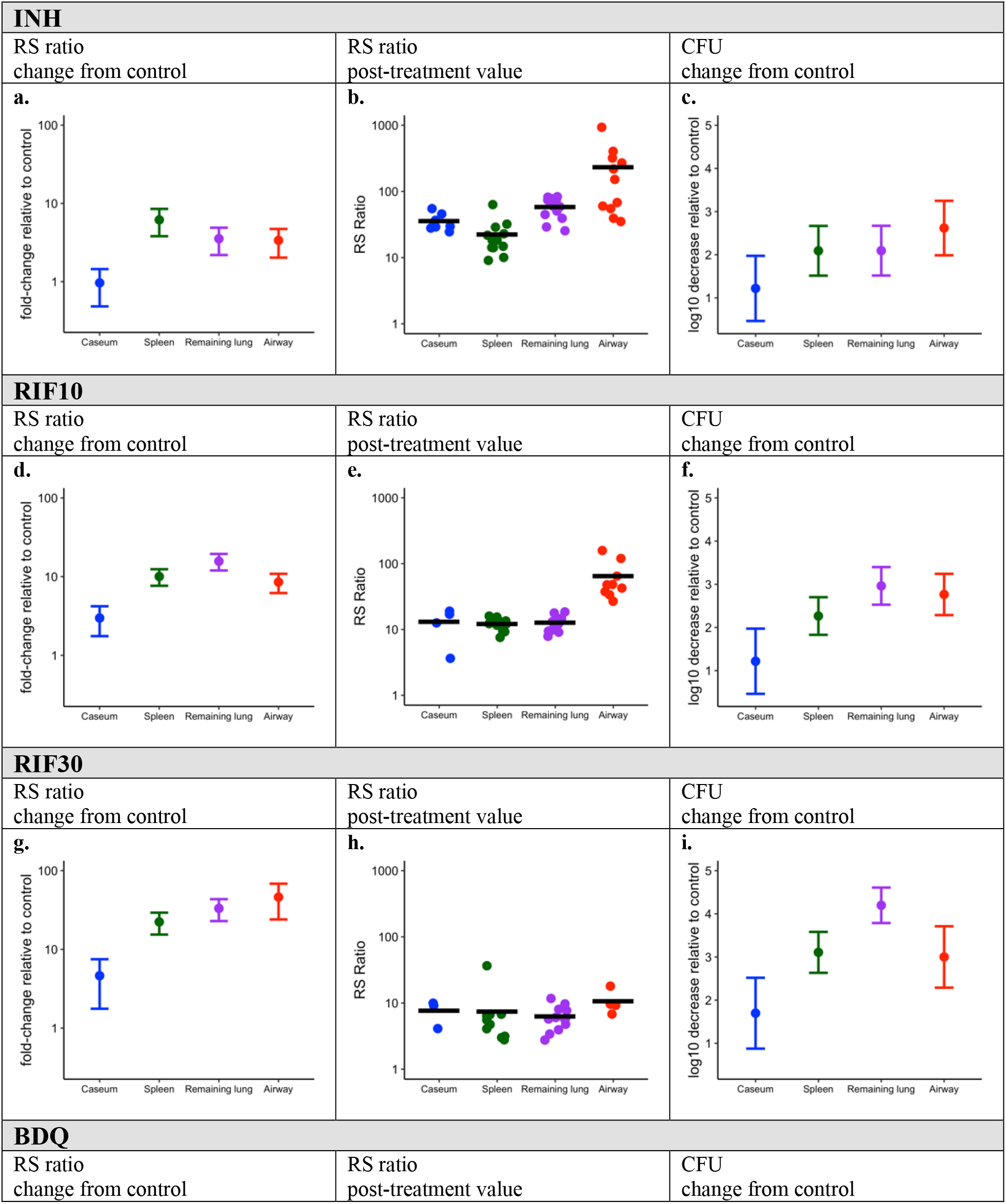
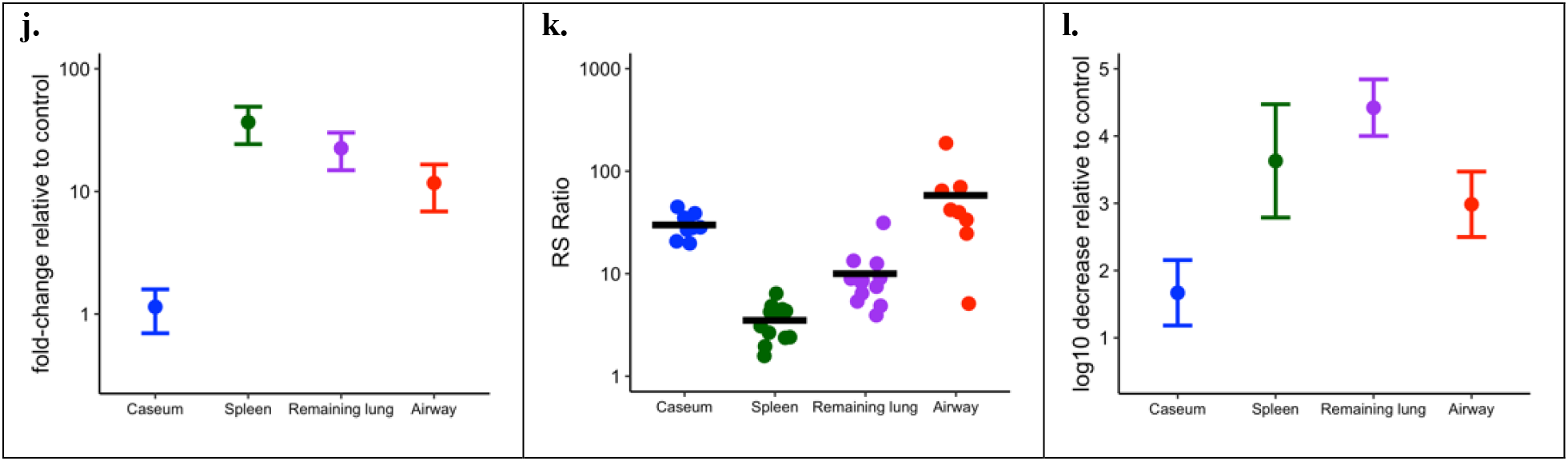
Effect of individual drugs and doses on the RS ratio and CFU in four microenvironments of the C3HeB/FeJ mouse. The fold change in RS ratio relative to untreated control in four microenvironments is shown for INH (**a**), RIF10 (**d)**, RIF30 (**g**) and BDQ (**j**). The central dot indicates the average fold-decrease in the RS ratio to control. Whiskers represent the 95% confidence interval. The absolute RS ratio value following treatment is shown for INH (**b**), RIF10 (**e**), RIF30 (**h**) and BDQ (**k**). Dots indicate values from individual mice. Horizontal lines indicate group means. The change in CFU relative to control is shown for INH (**c**), RIF10 (**f**), RIF30 (**i**) and BDQ (**l**). The central dot indicates the average log10 decrease in CFU relative to control. Whiskers represent the 95%confidence interval. Caseum, spleen, remaining lung and airway are represented in blue, green, purple and red, respectively.

### Effect of INH in caseum, spleen, remaining lung and airway

As noted above, INH caused minimal change in the RS ratio relative to control in caseum (**Fig 4a**). By contrast, INH reduced the RS ratio 5.6-fold in spleen, 3.5-fold in remaining lung and 3.0-fold in airway, representing a significantly greater reduction in all microenvironments other than in caseum (minimum *P*=0.01). Following treatment with INH, the absolute value of the RS ratio did not differ significantly between caseum and spleen (*P*= 0.3) or remaining lung (*P*=0.5) (**Fig 4b**, Supplemental Table 4). Following treatment, the RS ratio remained significantly higher in airway than in any other microenvironment (minimum *P*=0.01). The log10 reduction in CFU relative to control was statistically indistinguishable between microenvironments, with the exception that CFU declined marginally more in airway than in caseum (*P*=0.04) (**Fig 4c**).

### Effect of RIF10 in caseum, spleen, remaining lung and airway

RIF10 reduced the RS ratio 2.6-fold relative to control in caseum (**Fig 4d**). By contrast, RIF10 reduced the RS ratio 9.2-fold in spleen, 15.5-fold in remaining lung and 7.7-fold in airway, representing a significantly greater reduction in all other microenvironments than in caseum (minimum *P*=0.01). Following treatment with RIF10, the absolute value of RS ratio was similar in caseum, spleen and remaining lung but remained significantly higher in airway (minimum *P*-value <0.00001) (**Fig 4e**, Supplemental Table 4). RIF10 reduced CFU to statistically indistinguishable degrees in spleen, remaining lung and airway (2.0 to 2.4 log_10_ CFU) (**Fig 4f**). RIF10 reduced CFU significantly less in caseum than in remaining lung (*P*=0.002) or airway (*P*=0.009).

### Effect of RIF30 in caseum, spleen, remaining lung and airway

RIF30 reduced the RS ratio 4.1-fold relative to control in caseum (**Fig 4g**). By contrast, RIF30 reduced the RS ratio 20.4-fold in spleen, 32.6-fold in remaining lung and 41.7-fold in airway, representing a significantly greater reduction in all other microenvironments than in caseum (minimum *P*=0.006). Following treatment with RIF30, the absolute value of the RS ratio was similar in all microenvironments (**Fig 4h**, Supplemental Table 4). RIF30 reduced CFU significantly less in caseum than in spleen (*P*=0.04) or remaining lung (*P*=0.0001). Evaluation of exposure-response showed that RIF30 reduced the RS ratio significantly more than RIF10 in spleen, remaining lung and airway (minimum *P*=0.004). RIF30 also reduced CFU significantly more in spleen and remaining lung than in caseum (minimum *P*=0.002). For caseum, there was no significant difference between RIF30 and RIF10 for either the RS ratio or CFU.

### Effect of BDQ in caseum, spleen, remaining lung and airway

BDQ had minimal effect on the RS ratio in caseum (**Fig 3j**). By contrast, BDQ reduced the RS ratio 33.5-fold in spleen, 22.1-fold in remaining lung and 10.6-fold in airway, representing a significantly greater reduction in these microenvironments than in caseum (minimum *P=*0.0008). Following treatment with BDQ, the absolute value of RS ratio remained significantly higher in caseum than in spleen (*P*<0.00001) or remaining lung (*P*=0.0003) (**Fig 3k**). BDQ reduced CFU significantly less in caseum than in any other microenvironment (minimum *P*=0.004) (**Fig 3j**).

### Drug distribution within microenvironments

To determine how standard potency metrics compare to concentrations achieved in each lung microenvironment and help interpret the efficacy results, we applied drug quantitation by LC/MS-MS of samples collected by laser-capture microdissection (LCM) in thin lesion sections. Quanitification of LCM samples after 17 doses demonstrated that RIF10 was present at concentrations well above the serum shifted MIC (fMIC) (Supplemental Table 5) in all sampled tissue compartments including caseum. Trough samping 24 hours after dosing showed unchanged RIF concentration in caseum (**Fig 5a**). At day 17, INH was present at concentrations well above fMIC in all sampled tissue compartments including caseum one hour after dosing but rapidly exited, falling below fMIC after 24 hours (**Fig 5b**). At day 17, BDQ concentrations were similar at peak and trough and displayed a gradient in which concentrations observed in inner caseum were 16-to 28-fold lower than in the outer caseum, which showed to be proportional to distance from the edge of the granuloma (**Fig 5c, e-f**). The BDQ gradient was much more pronounced with a single dose of BDQ, wherein found levels of BDQ ranged from 1,490 ng/g (peak) to 2,465 ng/g (trough) in outer caseum to below the limit of quantification in inner caseum (**Fig 5d**).

**Fig 5.**
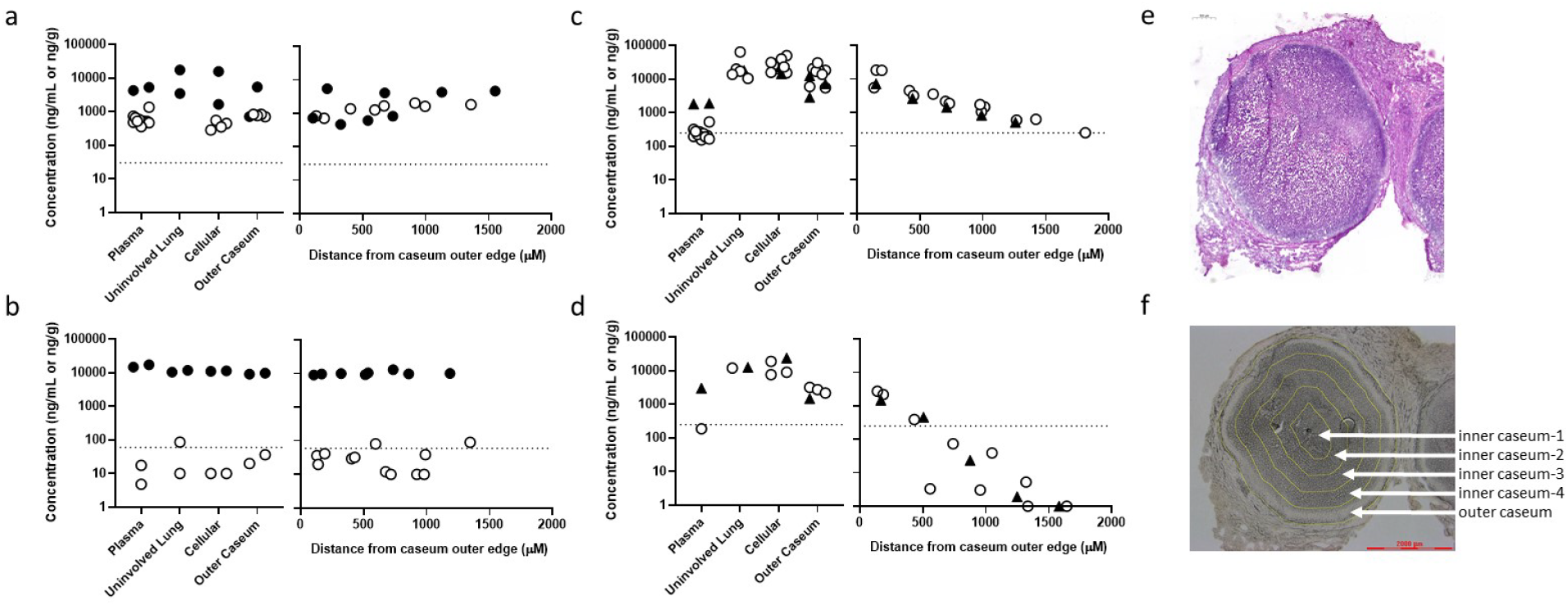
Drug distribution from plasma into different lung compartments and within caseum. (a-c) Concentration of RIF10 (a), INH (b) and BDQ (c) in caseum after 17 doses. Samples were collected from plasma and lung sections and at varying distances from the caseum outer edge at 1 hour (filled circles), 5 hours (filled triangles), or 24 hours (open circles) after dosing. Dotted line indicates the serum shifted MIC for each drug. (d) Concentration of BDQ in caseum after a single dose, displayed with the same formatting as in a-c. (e) Hematoxylin and eosin staining of Type I granuloma in C3HeB/FeJ mouse. (f) Regions of the caseum sampled by laser-capture microdissection.

## DISCUSSION

Application of the RS ratio and CFU burden demonstrated that lung microenvironments of the C3HeB/FeJ mouse harbor distinct *Mtb* phenotypes that have markedly different degrees of ongoing rRNA synthesis. Microenvironmental phenotypes appeared to be a determinant of drug effectiveness. All drugs had the lowest activity against *Mtb* populations in caseum. Even when tissue drug concentrations were above the serum shifted MIC, certain drugs were more effective against certain microenvironmental *Mtb* populations than others. Our *in vivo* observations in C3HeB/FeJ mice support and extend results showing higher drug tolerance and differential drug activity against *Mtb* in explanted rabbit caseum *ex vivo^3,23^* These results highlight the importance of considering PK and PD jointly. Combining a PD readout of a fundamental cellular process with lesional PK data may help to identify drugs and combinations that optimally target *Mtb* populations in caseum, a critical obstacle to shorter TB treatments.

Our findings are consistent with the long-standing hypothesis that one reason TB requires prolonged therapy is that unique tissue microenvironments harbor “special populations” that vary in their ability to withstand drug exposure.^1,7^ The RS ratio showed that *Mtb* maintains low ongoing rRNA synthesis in caseum, indicating a quiescent slowly-replicating population. This is consistent with the *ex vivo* rabbit caseum model in which which CFU and chromosomal equivalents showed no evidence of*Mtb* replication.^3^ Indeed, our RS ratio results indicated a similar low level of rRNA synthesis for *Mtb* in caseum *in vivo* and to caseum surrogate *ex vivo*, which in the absence of drug-treatments, is indicative of a dramatically reduced replication rate. Conversely, the *Mtb* populations of other environments appear to be replicating more quickly, with highest RS ratios in the oxygen-rich airway.

PD markers play a central role in drug evaluation because they are the readout used to assess drug effect. The historical standard PD marker is enumeration of the burden of *Mtb* capable of growth on solid culture (*i.e*., CFU).^24,25^ The RS ratio introduces a fundamentally different type of readout that measures a central bacterial cellular process rather than burden. As highlighted below, this new readout of a physiologic property revealed differences in drug effects that were not discernable based on CFU, thereby providing a new perspective on the effect of individual drugs against different microenvironmental phenotypes.

INH illustrates that achieving a concentration that exceeds the fMIC is insufficient if a microenvironment harbors an *Mtb* phenotype that is capable of withstanding a drug’s mechanism of action. For the quiescent *Mtb* population of caseum that likely has a low need for ongoing mycolic acid synthesis, effective concentrations of INH had no effect on the RS ratio. By contrast, for the actively replicating *Mtb* populations in spleen, remaining lung, and airway, interruption of new cell wall synthesis by INH reduced the RS ratio 3.0-5.6-fold. This is consistent with a body of evidence showing that INH is effective primarily against replicating *Mtb*, with minimal activity against non-replicating *Mtb*.^3,26,27^ In contrast to the RS ratio, CFU did not reveal differences between the effect of INH in different microenvironments, with the exception that CFU declined marginally more in airway than caseum. INH highlights the importance of bacterial phenotype and the information provided by a new molecular marker of rRNA synthesis.

RIF reduced the RS ratio in all sampled tissue compartments, indicating that RIF has activity against the phenotypes found in all microenvironments. Comparison of RIF10 and RIF30 demonstrated a strong dose-response relationship in spleen, remaining lung and airway.

Nonetheless both RS ratio and CFU indicated that RIF was less effective in caseum than in any other microenvironment. This highlights that caseum phenotypes are better able to tolerate even high concentrations of RIF, the prototypical sterilizing agent that is the backbone of existing front-line TB treatment.

BDQ represents a different example. In microenvironments that achieved high BDQ concentrations (*e.g*., remaining lung), BDQ matched the potency of RIF30 as measured by RS ratio and CFU. By contrast, drug distribution of BDQ into caseum appeared to be proportional to number of doses administered and inversely correlated with distance from the outer caseum edge. Spatial drug distribution studies by LCM LC/MS-MS suggested that after 2.5 weeks of treatment the concentration of BDQ ranged from 14.6 μg/g in outer caseum to as low as 0.25 μg/g in inner caseum and the effect on RS ratio was correspondingly negligible. Comparing the concentration gradient after 1 and 17 doses (**Fig. 5c-d**) suggests that BDQ slowly penetrates avascular caseum and may not have reached steady state at the 2.5 week mark, consistent with the slow onset of efficacy in early bactericidal activity Phase II trial.^28^ The current results do not indicate what the effect of BDQ might be were adequate caseum concentrations achieved.

Our observation that all drugs have diminished effectiveness in C3HeB/FeJ mouse caseum is broadly consistent with previous results from the rabbit *ex vivo* caseum model.^3^ Using CFU as a readout, Sarathy *et.al*., found that the caseum MBC of first and second line drugs was higher in rabbit caseum than under standard *in vitro* conditions with replicating cultures. Because the RS ratio is a direct molecular PD readout that does not require confounding steps such as recovery and outgrowth of bacteria, the RS ratio enabled this first comparison of phenotypes and drug effect across tissue microenvironments. The RS ratio extends and complements results from the *ex vivo* rabbit caseum model by measuring a fundamental cellular process rather than bacillary burden.

In contrast to our results, a recent manuscript that evaluated a fluorescent reporter strain in C3HeB/FeJ mouse lesions reported that *Mtb* was more actively replicating in the caseum core than in the cuff and the *Mtb* population of the core showed greater susceptibility to INH.^29^ Importantly, the previous report evaluated a different timepoint than tested here. Lavin and Tan^29^ evaluated mice 6 weeks post infection (a time point at which previous studies would suggest lesions are in the maturation process^19^) and identified rapidly replicating caseum phenotypes that were highly susceptible to INH. By contrast, we evaluated mice 12.5 weeks post-infection (a time point at which lesions are mature and well-encapsulated with high *Mtb* burden) and identified slowly replicating caseum phenotypes that were INH-tolerant. In combination, these observations appear to highlight how the evolution of lesions and phenotypes over time dictates drug effectiveness. Time-course studies using a single consistent assay method are needed to confirm this hypothesis.

This report has several limitations. This proof-of-concept study included limited PK measurements that were insufficient for statistical pharmacokinetic modeling. Future studies can maximize the value of lesional analysis by pairing lesional PD (based on RS ratio and CFU) with comprehensive lesional PK profiles. Second, the RS ratio provides a snapshot of average rRNA synthesis across an entire *Mtb* population. Using *in situ* hybridization with single-bacillary resolution, we have previously shown that caseum harbors multiple *Mtb* phenotypes with differing degrees of ongoing rRNA synthesis.^8,15^ In future studies, the joint PK/PD evaluation demonstrated here can be extended with even more spatially granular methods.

This study has shown that different tissue microenvironments harbor phenotypically distinct *Mtb* populations that respond to drug treatment differently. As a molecular readout of a fundamental cellular process, the RS ratio provides a new metric for evaluating efficacy in lesions that does not rely on bacillary burden. This report points to a new era in which lesional PK can be combined with molecular assays of bacterial cellular processes like the RS ratio to design regimens that optimally target caseum phenotypes and achieve the end-goal of shorter, more effective TB treatments.

## MATERIALS AND METHODS

### Drug susceptibility profiling

The Minimal Inhibitory Concentration (MIC) was determined for RIF, INH, and BDQ against *Mtb* Erdman in 7H9 media supplemented with 0.2% [v:v] glycerol and 10% [v:v] ADC, with 0.05% [v:v] Tween-80 (7H9 media). MICs were determined by a broth microdilution assay using two-fold serial drug dilutions. The lowest consecutive antimicrobial concentration that showed a ≥ 80% reduction in OD600 relative to drug-free control wells, was regarded as the MIC. In parallel, the MICs were also determined in presence of 4% (w/v) human serum albumin (Sigma # A1653) which is defined here as the serum shifted MIC.

### Animals

Female specific pathogen-free C3HeB/FeJ mice, age 8-10 weeks, were purchased from Jackson Laboratories (Bar Harbor, ME). Mice were housed in an animal bio-safety level III (ABSL-3) facility employing autoclaved bedding, water and mouse chow *ad libitum*. All procedures were approved by the Colorado State University Institutional Animal Care and Use Committee (IACUC) (Reference numbers of approved protocol: KP 1515).

### Aerosol infection

C3HeB/FeJ mice were exposed to a low dose aerosol infection with the *Mtb* Erdman strain (TMCC 107) using a Glas-Col inhalation exposure system resulting in an average of 50-75 bacteria in lungs one day following aerosol.^8,30^ Infected mice were observed and weighed at least once a week. Starting at day 21 until the start of therapy, mice were observed and weighed two to three times per week, due to the increased incidence of morbidity and mortality associated with clinical TB disease. Any mice exhibiting clinical symptoms of illness were humanely euthanized.

### Drug preparation and drug treatment

Rifampin (Sigma) and isoniazid (Sigma) were dissolved in sterile water. Bedaquiline fumarate (PharmaBlock) was dissolved in 20% (w/v) 2-HPCD and formulated as described previously.^31,32^ For drug efficacy experiments, C3HeB/FeJ mice were randomized (n=12 per group) and dosed once daily starting 10 weeks post aerosol with rifampin at 10 mg/kg or 30 mg/kg, isoniazid at 25 mg/kg, or bedaquiline fumarate at 25 mg/kg, 7 days per week by oral gavage in 0.2 mL volume for the total number of doses indicated.

### Histological analysis

A separate group of mice were sacrificed and whole lungs fixed for histopathology purposes. Lung perfusions were performed by clamping the caudal vena cava with straight tipped hemostats and cutting the vena cava between the hemostats and the liver to allow blood to drain. A 24 guage 0.75 inch catheter was inserted into the right ventricle of the heart and blood flushed using PBS with 0.04% [w:v] EDTA followed by 10 mL of 4% paraformaldehyde (PFA). Lungs were recovered and placed into a histology cassette and incubated for 48 h in 4% PFA for 48 hours prior to transfer to PBS. Lungs were sectioned and stained with hematoxylin-eosin.

### Collection of samples for bacterial enumeration and RS ratio

Each mouse was individually euthanized by CO_2_ narcosis followed by cardiac puncture. Airway was sampled by bronchial alveolar lavage (BAL) by passing a total of 2.5 mL PBS into lungs through a 24 guage 0.75 inch catherter inserted into the trachea. 0.5 mL of BAL was used for CFU enumeration. 0.5 mL was placed into 2 mL CAMM-RPS buffer to preserve RNA. Remaining BAL was collected by cytospin for Wright’s giemsa staining. Spleens were aseptically collected. Lungs were recovered, photographed and diagramed. Lung lobes featuring large encapsulated caseous necrotic granulomas (Type I) were used to dissect caseum samples. All tissue samples were divided in half and weights were recorded. Samples were flash frozen in liquid nitrogen and stored at −80°C prior to further processing.

Tissues were disrupted with a tissue homogenizer (Precellys, Bertin Instruments, Rockville, MD) in PBS plus 10% bovine serum albumin for CFU enumeration on 7H11 agar further supplemented with 0.4% activated charcoal to prevent drug carry-over as described previously.^15^ Colonies were enumerated after at least 28 days of incubation at 37 °C and plates were incubated for 8 weeks to ensure all viable colonies were detected. For RNA extraction, tissues were homogenized in Trizol.^15^

### Caseum surrogate model

PMA-differentiated THP-1 macrophages were infected with irradiated *Mtb* (BEI Resources) at an approximate MOI of 1:50. Foamy macrophages were washed three times with PBS, followed by three freeze–thaw cycles to lyse the cells and incubation at 75 °C for 20 min to denature proteins in the matrix. The caseum surrogate matrix was rested for 3 days at 37 °C and stored at −20 °C prior to inoculation.

### Collection of samples from caseum surrogate

*Mtb* HN878 grown to mid log phase in 7H9 media were harvested by centrifugation, suspended in sterile water, and inoculated into caseum surrogate^21,22^ at 10^8^ CFU/mL. The inoculated caseum surrogate was mixed briefly using 1.4 mm zirconium oxide beads (CK14 soft tissue; Precellys, Bertin Instruments, Rockville, MD) to achieve a homogenious suspension. After 8 weeks incubation in a sealed tube at 37°C to allow for physiologic and metabolic adaptation to the lipid-rich environment, CAMM-RPS buffer was added at 3× the total caseum surrogate volume to preserve RNA.

### Collection of samples for pharmacokinetic analysis

For the PK experiments, 9 to 10 C3HeB/FeJ mice were dosed starting 10 weeks post aerosol as above. Treatment occurred 7 days per week for up to a total of 17 doses. Drugs were prepared as described above for the efficacy studies. Mice were euthanized, and plasma and tissues were collected at two time points, selected based on the plasma C_max_ (1 or 5 hours) and C_min_ (24 hours). Whole blood was obtained via cardiac puncture and processed in plasma separator tubes (Becton, Dickinson and Co., Franklin Lakes, NJ) centrifuged at 3,000 RCF for 2 min at 4°C, aliquoted into Eppendorf microcentrifuge tubes and stored at −80°C until analysis. Mice with pronounced lung pathology were selected to collect samples for spatial drug quantitation by gravity assisted LCM. Briefly, whole lung samples were collected on clear disposable base molds (Fisher Scientific, Hampton, NH, USA). Using forceps, tissues collected on trays placed on a pre-chilled aluminum block held in liquid nitrogen, and frozen within 1-2 minutes. Tissue trays containing frozen lobes were wrapped in foil squares, placed individually into labeled zip-lock bags and immediately transferred onto dry ice.

### Drug quantitation by HPLC coupled to tandem mass spectrometry (LC/MS-MS)

Drug levels in tissue were determined by spatial quantitation in thin tissue sections by LCM followed by LC/MS-MS analysis of microdissected areas.^33^ The benefit of the LCM approach is the ability to obtain absolute drug levels in defined areas of Type I lung lesions such as caseum (the core of the caseous necrotic lesion) without cross-contamination. 25 μm-thick tissue sections were cut from infected mouse lung biopsies using a Leica CM 1860UV (Buffalo Grove, IL) and thaw-mounted onto 1.4 μm-thick Leica PET-Membrane FrameSlides (Buffalo Grove, IL). Adjacent 10 μm thick tissue sections were thaw-mounted onto standard glass microscopy slides for H&E staining. Sequential rings of necrotic tissue proceeding from the border with the cellular rim to core of the caseous compartment were demarcated for each tissue section by creating a mask of caseum and processing it using the Exact Euclidean Distance Transform plugin in ImageJ (National Institutes of Health, MD). Sequential rings of necrotic tissue were then dissected using a Leica LMD6 system (Buffalo Grove, IL). Dissected lesion tissues were collected into 0.25 mL standard PCR tubes and immediately transferred to the −80°C.

RIF, BDQ, INH and verapamil (VER) were purchased from Sigma Aldrich (St. Louis, MO); RIF-d8, BDQ-d6 and INH-d4 were purchased from Toronto Research Chemicals (Ontario). Drug free lung and K2EDTA plasma from CD-1 mice was obtained from BioIVT (Westbury, NY) for use as blank matrices to build standard curves. Neat 1 mg/mL DMSO stocks were serial diluted in 50:50 acetonitrile/water to create standard curve and quality control spiking solutions. 10 μl of neat spiking solutions were added to 2 μl of lesion homogenate prior to extraction. On the day of analysis, samples were extracted by adding 50 μl of extraction solution (acetonitrile/methanol, 1:1) containing 1, 10, 4 and 5 ng/mL of VER, INH-d4, RIF-d8 and BDQ-d6 respectively. Extracts were sonicated for 5 minutes and centrifuged at 10,000 rpm for 5 minutes. 40 μl of supernatant was transferred to 96-well deep plates for LC-MS/MS analysis. For INH quantitation, 40 μl of 2% cinnamaldehyde in methanol was added to derivatize isoniazid prior to analysis. For RIF quantitation, 5 μl of 75 mg/mL ascorbic acid was added to extracts to stabilize RIF during analysis. LC/MS-MS analysis was performed on a Sciex Qtrap 6500+ triple-quadrupole mass spectrometer coupled to a Shimadzu Nexera X2 UHPLC system. Chromatography was performed on an Agilent Zorbax SB-C8 column (2.1×30mm; particle size 3.5 μm) using a reverse phase gradient. Deionized water with 0.1% formic acid (FA) was used for the aqueous mobile phase and 0.1% FA in acetonitrile for the organic mobile phase. Multiple-reaction monitoring (MRM) of precursor/fragment transitions in electrospray positive-ionization mode was used to quantify the analytes. MRM transitions of 823.30/791.30, 555.00/58.00, 252.200/80.30 455.40/165.20, 831.30/799.40, 561.00/64.00 and 256.20/84.30 were used for RIF, BDQ, INH, VER, RIF-d8, BDQ-d6 and INH-d4 respectively. Data processing was performed using Analyst software version 1.6.3 (Sciex).

### RNA extraction and RS ratio profiling

RNA was extracted from tissue and BAL samples after homogenization in Trizol.^15^ Briefly, eukaryotic cellular debris was pelleted and the lysate was mixed with chloroform. A solution of 50% isopropanol, 0.8M sodium citrate, and 1.2M sodium chloride was mixed with the aqueous phase and nucleic acids were precipitated overnight at 4°C. Nucleic acids were pelleted by centrifugation, washed twice with 70% ethanol, and resuspended in water. DNA was digested in a multi-step process, with the initial digestion performed using two additions of Promega RQ1 Dnase followed by a 15 minute incubation at 37°C after each Dnase addition. RNA was further purified and DNA digested on a Maxwell RSC instrument (Promega) with the Maxwell RSC simplyRNA tissue kit. Purification was performed following the manufacturer’s instructions, with Dnase added at twice the recommended amount.

Purified RNA was reverse transcribed to cDNA using SuperScript III VILO cDNA synthesis kit (Invitrogen) at 42°C for 120 minutes. The RS ratio was measured by quantification of ETS1 and 23S rRNA gene transcripts in the cDNA using droplet digital PCR in a duplexed reaction. Droplet digital PCR reactions were performed using the QX200 Droplet Digital PCR system with ddPCR SuperMix for Probes (no dUTP) (Bio-Rad). The RS ratio was calculated from each duplexed reaction by the QX200 Droplet Digital PCR system software (QuantaSoft AP, Bio-Rad).

## Statistical analysis

A one-way ANOVA was performed to compare the effect of different drugs or microenvironments on CFU and the RS ratio followed by a multiple comparison analysis of variance using a one-way Tukey test. Differences were considered significant at the 95% confidence level (*P*< 0.05). Data management, plotting and post-modeling analysis were conducted using R (v 4.2.1; R Development Core Team, Vienna, Austria).

## SUPPLEMENTAL MATERIAL

Supplemental material is available online only.

## ACKNOWLEDGMENTS

NDW acknowledges funding from Veterans Affairs (1I01BX004527-01A1) and the US National Institutes of Health (1R01AI127300-01A1). NDW and MIV acknowledge funding from the US National Institutes of Health (1R21AI135652-01). NDW, MIV, GTR, and RMS acknowledge funding from the Bill & Melinda Gates Foundation (OPP1213947). GTR, NDW, and VD acknowledge funding from the Bill & Melinda Gates Foundation (OPP1126594).

## AUTHOR CONTRIBUTIONS

NDW, GTR, RMS, MIV and BKP conceived of the project and planned the experiments. NDW and GTR drafted the manuscript. AAB, MER, and LMM conducted animal experiments. KR, SP, and RAM performed molecular assays. FK and JPS performed PK analyses. CDA, MDZ, JPE, VD analyzed molecular and PK results.

## Notes

### Competing Interest Statement

The authors have declared no competing interest.

