## supplemental tables for "Lung microenvironments harbor *Mycobacterium tuberculosis* phenotypes with distinct treatment responses"

**Affiliation:**

† Corresponding author

### 1. Supplemental Results

#### 1.1. Supplemental Table 1

**Supplemental Table 1.** Pairwise comparisons of CFU and RS ratio across microenvironments in C3HeB/FeJ mice prior to drug treatment. The mean difference between CFU or mean fold-change between RS ratio in the comparator microenvironment versus the reference microenvironment is shown with associated *P*-value. A positive difference indicates that CFU or the RS ratio was higher in the comparator than in the reference. A fold-change <1 indicates that the RS ratio was lower in the comparator than in the reference. *P*-values <0.05 are highlighted in bold.

| Comparator | Reference | CFU | | RS ratio | |
|---|---|---|---|---|---|
| | | **$Log_{10}$ difference** | ***P*-value** | **Fold-change** | ***P*-value** |
| Caseum | Spleen | 3.2 | **<0.00001** | -3.7 | **<0.00001** |
| Caseum | Remaining Lung | 2.4 | **<0.00001** | -6.4 | **<0.00001** |
| Caseum | Airway | 3.1 | **<0.00001** | -14.5 | **<0.00001** |
| Remaining Lung | Spleen | 0.8 | **0.02** | 1.7 | **0.01** |
| Remaining Lung | Airway | 0.6 | 0.1 | -2.2 | **0.0003** |
| Spleen | Airway | -0.2 | 0.9 | -3.8 | **<0.00001** |

#### 1.2. Supplemental Table 2

**Supplemental Table 2.** Pairwise comparisons of CFU and RS ratio for individual drugs within specific microenvironments. The mean difference between CFU or mean fold-change between RS ratio for each drug versus the reference is shown with associated *P*-value. A positive difference indicates that CFU or the RS ratio was higher for the comparator drug than in the reference. A fold-change <1 indicates that the RS ratio was lower in the comparator than in the reference. *P*-values <0.05 are shown in bold.

| Comparator | Reference | CFU | | RS ratio | |
|---|---|---|---|---|---|
| | | **Mean $\log_{10}$ difference** | ***P*-value** | **Mean fold-change** | ***P*-value** |
| **Caseum** | | | | | |
| INH | CONTROL | -1.00 | **0.02** | 1.19 | 1.0 |
| RIF10 | CONTROL | -1.00 | **0.06** | -2.61 | **0.04** |
| RIF30 | CONTROL | -1.48 | **0.005** | -4.05 | **0.003** |
| BDQ | CONTROL | -1.45 | **0.0003** | -1.00 | 1.0 |
| RIF10 | INH | 0.00 | 1.0 | -3.09 | **0.005** |
| RIF30 | INH | -0.48 | 0.7 | -4.80 | **0.0004** |
| BDQ | INH | -0.45 | 0.4 | -1.19 | 1.0 |
| RIF30 | RIF10 | -0.48 | 0.7 | -1.55 | 0.7 |
| BDQ | RIF10 | -0.45 | 0.6 | 2.61 | **0.02** |
| BDQ | RIF30 | 0.03 | 1.0 | 4.05 | **0.001** |
| **Spleen** | | | | | |
| INH | CONTROL | -1.85 | **<0.00001** | -5.63 | **<0.00001** |
| RIF10 | CONTROL | -2.03 | **<0.00001** | -9.18 | **<0.00001** |
| RIF30 | CONTROL | -2.87 | **<0.00001** | -20.43 | **<0.00001** |
| BDQ | CONTROL | -3.39 | **<0.00001** | -33.53 | **<0.00001** |
| RIF10 | INH | -0.17 | 0.8 | -1.63 | 0.1 |
| RIF30 | INH | -1.02 | **<0.00001** | -3.63 | **<0.00001** |
| BDQ | INH | -1.54 | **<0.00001** | -5.95 | **<0.00001** |
| RIF30 | RIF10 | -0.84 | **0.00005** | -2.22 | **0.002** |
| BDQ | RIF10 | -1.37 | **<0.00001** | -3.65 | **<0.00001** |
| BDQ | RIF30 | -0.52 | 0.2 | -1.64 | 0.1 |
| **Remaining Lung** | | | | | |
| INH | CONTROL | -1.50 | **0.0002** | -3.48 | **<0.00001** |
| RIF10 | CONTROL | -2.37 | **<0.00001** | -15.46 | **<0.00001** |
| RIF30 | CONTROL | -3.60 | **<0.00001** | -32.63 | **<0.00001** |
| BDQ | CONTROL | -3.83 | **<0.00001** | -22.13 | **<0.00001** |
| RIF10 | INH | -0.87 | 0.05 | -4.44 | **<0.00001** |
| RIF30 | INH | -2.10 | **<0.00001** | -9.37 | **<0.00001** |
| BDQ | INH | -2.33 | **<0.00001** | -6.35 | **<0.00001** |
| RIF30 | RIF10 | -1.23 | **0.002** | -2.11 | **0.0002** |
| BDQ | RIF10 | -1.46 | **0.0002** | -1.43 | 0.2 |
| BDQ | RIF30 | -0.22 | 1.0 | 1.47 | 0.1 |
| **Airway** | | | | | |
| INH | CONTROL | -2.01 | **0.003** | -3.04 | **0.01** |
| RIF10 | CONTROL | -2.16 | **0.001** | -7.70 | **<0.00001** |
| RIF30 | CONTROL | -2.40 | **0.01** | -41.65 | **<0.00001** |
| BDQ | CONTROL | -2.38 | **0.0005** | -10.57 | **<0.00001** |
| RIF10 | INH | -0.15 | 1.0 | -2.53 | 0.09 |
| RIF30 | INH | -0.38 | 1.0 | -13.69 | **<0.00001** |
| BDQ | INH | -0.37 | 1.0 | -3.47 | **0.01** |
| RIF30 | RIF10 | -0.24 | 1.0 | -5.41 | **0.004** |
| BDQ | RIF10 | -0.22 | 1.0 | -1.37 | 0.9 |
| BDQ | RIF30 | 0.02 | 1.0 | 3.94 | **0.03** |

#### 1.3. Supplemental Table 3

**Supplemental Table 3.** Pairwise comparisons of reduction in CFU and RS ratio from control across microenvironments for individual drugs. The mean difference between CFU or mean fold-change between RS ratio in the comparator microenvironment versus the reference microenvironment is shown with associated *P*-value. A positive difference indicates that CFU or the RS ratio was reduced to a greater degree in the comparator than in the reference. A fold-change <1 indicates that the RS ratio was lower in the comparator than in the reference. *P*-values <0.05 are shown in bold.

| Comparator | Reference | CFU | | RS ratio | |
|---|---|---|---|---|---|
| | | Mean $\log_{10}$ difference | *P*-value | Mean fold-change | *P*-value |
| **INH** | | | | | |
| Caseum | Spleen | -0.87 | 0.3 | -5.19 | **0.0008** |
| Caseum | Remaining lung | -0.87 | 0.3 | -2.57 | **0.006** |
| Caseum | Airway | -1.40 | **0.04** | -2.41 | **0.01** |
| Remaining lung | Spleen | 0.00 | 1.0 | -2.62 | 0.2 |
| Remaining lung | Airway | -0.52 | 0.6 | 0.17 | 1.0 |
| Spleen | Airway | -0.53 | 0.6 | 2.79 | 0.2 |
| **RIF10** | | | | | |
| Caseum | Spleen | -1.05 | 0.1 | -7.06 | **0.00007** |
| Caseum | Remaining lung | -1.75 | **0.002** | -12.72 | **<0.00001** |
| Caseum | Airway | -1.55 | **0.009** | -5.55 | **0.001** |
| Remaining lung | Spleen | 0.70 | 0.1 | 5.66 | 0.08 |
| Remaining lung | Airway | 0.20 | 0.9 | 7.18 | **0.02** |
| Spleen | Airway | -0.50 | 0.4 | 1.51 | 0.8 |
| **RIF30** | | | | | |
| Caseum | Spleen | -1.41 | **0.04** | -17.71 | **0.0004** |
| Caseum | Remaining lung | -2.50 | **0.0001** | -28.52 | **0.00008** |
| Caseum | Airway | -1.30 | 0.1 | -41.51 | **0.006** |
| Remaining lung | Spleen | 1.09 | **0.01** | 10.81 | 0.3 |
| Remaining lung | Airway | 1.20 | **0.04** | -12.99 | 0.7 |
| Spleen | Airway | 0.11 | 1.0 | -23.80 | 0.2 |
| **BDQ** | | | | | |
| Caseum | Spleen | -1.96 | **0.002** | -35.52 | **0.00001** |
| Caseum | Remaining lung | -2.75 | **<0.00001** | -21.34 | **0.00002** |
| Caseum | Airway | -1.32 | **0.004** | -10.57 | **0.0008** |
| Remaining lung | Spleen | 0.79 | 0.4 | -14.18 | 0.2 |
| Remaining lung | Airway | 1.44 | **0.0008** | 10.77 | 0.1 |
| Spleen | Airway | 0.65 | 0.6 | 24.95 | **0.004** |

#### 1.4. Supplemental Table 4

**Supplemental Table 4.** *P*-values from pairwise comparisons of the absolute RS ratio value in key microenvironments following treatment.

| **Comparator** | **Reference** | ***P*-value** |
|---|---|---|
| **INH** | | |
| Caseum | Spleen | 0.4 |
| Caseum | Remaining lung | 0.5 |
| Caseum | Airway | **0.006** |
| Remaining lung | Spleen | **0.02** |
| Remaining lung | Airway | **0.04** |
| Spleen | Airway | **0.0009** |
| **RIF 10** | | |
| Caseum | Spleen | 1.0 |
| Caseum | Remaining lung | 1.0 |
| Caseum | Airway | **0.00003** |
| Remaining lung | Spleen | 1.0 |
| Remaining lung | Airway | **0.00003** |
| Spleen | Airway | **0.00002** |
| **RIF 30** | | |
| Caseum | Spleen | 0.9 |
| Caseum | Remaining lung | 0.9 |
| Caseum | Airway | 0.8 |
| Remaining lung | Spleen | 1.0 |
| Remaining lung | Airway | 0.4 |
| Spleen | Airway | 0.3 |
| **BDQ** | | |
| Caseum | Spleen | **0.0004** |
| Caseum | Remaining lung | **0.008** |
| Caseum | Airway | 0.7 |
| Remaining lung | Spleen | **0.009** |
| Remaining lung | Airway | **0.004** |
| Spleen | Airway | **0.0006** |

#### 1.5. Supplemental Table 5

**Supplemental Table 5.** Broth microdilution MIC assays conducted with *Mycobacterium tuberculosis* Erdman in 7H9 medium media supplemented with 0.2% [v:v] glycerol and 10% [v:v] ADC, with 0.05% [v:v] Tween-80 (7H9 media) plus or minus 4% [w:v] human serum albumin after 8 days incubation at 37°C. Antimicrobials tested and final test concentrations in mg/L are listed. MIC values are called as the first consecutive well showing greater than or equal to 80% growth inhibition (GI80) of the average OD at 600 nm of the no drug DMSO only control wells.

| **compound** | **MIC for Mtb Erdman in mg/L in 7H9-media with:** | |
|---|---|---|
| | **no supplement** | **4% human serum albumin** |
| rifampin | 0.008 | 0.03 |
| isoniazid | 0.03 | 0.06 |
| bedaquiline | 0.125 | 0.25 |
